# Epiretinal stimulation with local returns enhances selectivity at cellular resolution

**DOI:** 10.1101/275958

**Authors:** Victoria H. Fan, Lauren E. Grosberg, Sasidhar S. Madugula, Pawel Hottowy, Wladyslaw Dabrowski, Alexander Sher, Alan M. Litke, E.J. Chichilnisky

**Affiliations:** Departments of Neurosurgery, Ophthalmology, and Hansen Experimental Physics Laboratory, Stanford University, Stanford CA, USA.; AGH University of Science and Technology, Faculty of Physics and Applied Computer Science, 30-059 Krakow, Poland.; Santa Cruz Institute for Particle Physics, University of California, Santa Cruz, CA, USA

**Keywords:** retinal electrophysiology, retinal ganglion cells, epiretinal prosthesis, artificial retina, local return, single-cell resolution, selectivity

## Abstract

**Objective:** Epiretinal prostheses are designed to restore vision in people blinded by photoreceptor degenerative diseases, by directly activating retinal ganglion cells (RGCs) using an electrode array implanted on the retina. In present-day clinical devices, current spread from the stimulating electrode to a distant return electrode often results in the activation of many cells, potentially limiting the quality of artificial vision. In the laboratory, epiretinal activation of RGCs with cellular resolution has been demonstrated with small electrodes, but distant returns may still cause undesirable current spread. Here, the ability of local return stimulation to improve the selective activation of RGCs at cellular resolution was evaluated.

**Approach:** A custom multi-electrode array (512 electrodes, 10 μm diameter, 60 μm pitch) was used to simultaneously stimulate and record from RGCs in isolated primate retina. Stimulation near the RGC soma with a single electrode and a distant return was compared to stimulation in which the return was provided by six neighboring electrodes.

**Main results:** Local return stimulation enhanced the capability to activate cells near the central electrode (<30 μm) while avoiding cells farther away (>30 μm). This resulted in an improved ability to selectively activate ON and OFF cells, including cells encoding immediately adjacent regions in the visual field.

**Significance:** These results suggest that a device that restricts the electric field through local returns could optimize activation of neurons at cellular resolution, improving the quality of artificial vision.

**Novelty & Significance:** The effectiveness of local return stimulation for enhancing the electrical activation of retinal neurons was tested using high-density multi-electrode recording and stimulation in isolated macaque retina. The results suggest that local returns may reduce unwanted evoked activity and thus optimize the selectivity of stimulation at cellular resolution. Similar patterns could be implemented in a future high-resolution prosthesis to permit a more faithful replication of normal retinal activity for the treatment of incurable blindness.

## Introduction

An epiretinal prosthesis produces artificial vision in patients blinded by photoreceptor diseases such as retinitis pigmentosa, by passing current through electrodes in an array implanted on the retinal surface. Electrical stimulation causes the surviving retinal ganglion cells (RGCs) to fire and thus transmit artificial visual signals to the brain. Although these prostheses can produce visual percepts (Humayun et al., 1996; Rizzo et al., 2003; Hornig et al., 2017; see Weiland and Humayun, 2014; see Goetz and Palanker, 2016), the coarse and unnatural patterns of retinal activity caused by simultaneous activation of many neighboring cells of different types almost certainly limit the quality of artificial vision.

In principle, higher resolution stimulation is possible with effective use of spatially patterned stimulation. Several approaches have been proposed. For example, current steering, which involves passing current through several electrodes simultaneously in customized patterns, has been used in a range of neural structures from the cochlea (Townshend and White, 1987; Firszt et al., 2007) to deep brain nuclei (Martens et al., 2011) to the retina (Jepson et al., 2013; Matteucci et al., 2013; Dumm et al., 2014; see Bareket et al., 2017). Similarly, the use of local returns (rather than distant returns) to limit current spread without customization has been investigated using subretinal stimulation (Palanker et al., 2005; Palanker, 2014; Habib et al., 2013; Flores et al., 2016), suprachoroidal stimulation (Wong et al., 2009; Cicione et al., 2012; Sinclair et al., 2016), epiretinal stimulation (Abramian et al., 2011), non-biological experiments (Dommel et al., 2005), and simulations (Joarder et al., 2011).

Although such approaches could improve the spatial resolution of stimulation to a degree, a major challenge for epiretinal stimulation is the fact that the different types of RGCs are interspersed on the surface of the retina. Therefore, to create the correct patterns of activity in different RGC types requires stimulating at the resolution of individual cells, a goal that has only been demonstrated in isolated macaque retina *ex vivo*, and even in these conditions is not always achievable (Sekirnjak et al., 2006, 2008; Jepson et al., 2013, 2014a; Grosberg et al., 2017). Therefore, it would be valuable to understand how effectively spatially patterned electrical stimulation can sharpen spatial patterns of activation, specifically at the resolution of individual cells. Although the success of customized current steering has been shown in an individual example cell (Jepson et al., 2014a), it remains unclear whether simpler open-loop techniques like local return stimulation can also be effective, and whether the improvements are observed systematically in many cells and retinas rather than isolated examples.

Here, we test whether local returns can enhance single-cell activation, using large-scale high-density recording and stimulation from isolated macaque retina (Hottowy et al., 2012; Jepson et al., 2013, 2014b; Grosberg et al., 2017). We evaluate how effectively a given target cell can be activated without activating a nearby non-target cell, using both local and distant return stimulation. We demonstrate that when the target cell is near the stimulating electrode and the non-target cell is farther away, local returns can enhance the selective activation of the target cell over the non-target cell. This enhancement permits selective activation of different cells and cell types encoding immediately adjacent portions of the visual field. The results provide support for using local return stimulation to more faithfully reproduce natural RGC activity in future high-resolution epiretinal prostheses.

## Methods

### Experimental setup

A custom 512-electrode system (Hottowy et al., 2008, 2012, Grosberg et al., 2017) was used to stimulate and record from hundreds of RGCs in isolated rhesus macaque monkey (*Macaca mulatta*) retina. Retinas were obtained from terminally anesthetized animals euthanized in the course of experiments by other laboratories. Briefly, eyes were hemisected in room light following enucleation. The vitreous was then removed and the posterior portion of the eye containing the retina was kept in darkness in warm, oxygenated, bicarbonate buffered Ames’ solution (Sigma). Patches of retina ~3 mm on a side were isolated under infrared light, placed RGC side down on the multielectrode array, and superfused with Ames solution at 33 °C. Electrodes were 8-10 μm in diameter and arranged in a 16 × 32 isosceles triangular lattice with 60 μm spacing between adjacent electrodes (Litke et al., 2004). Electrodes were electroplated with platinum. Voltage recordings were band-pass filtered between 43 and 5,000 Hz and sampled at 20 kHz.

### Electrical stimulation

Electrical stimuli were provided on one or more channels while recording RGC activity from all channels. Two types of stimulation patterns were tested (Fig. 1). The distant return pattern consisted of a charge-balanced triphasic pulse on a single electrode, with a bath ground (platinum wire ring) ~1 cm away (Fig. 1B). The triphasic pulse was made up of anodal/cathodal/anodal phases with relative current amplitudes of 2:-3:1 and phase duration of 50 μs (150 μs total duration) (Fig. 1A). These parameters were chosen to minimize the electrical artifact (Hottowy et al., 2012; Jepson et al., 2013; Grosberg et al., 2017). The local return pattern consisted of the same central electrode current, with simultaneous current waveforms of opposite sign and ⅙ amplitude on the six immediately surrounding return electrodes (Fig. 1C).

**Figure 1.**
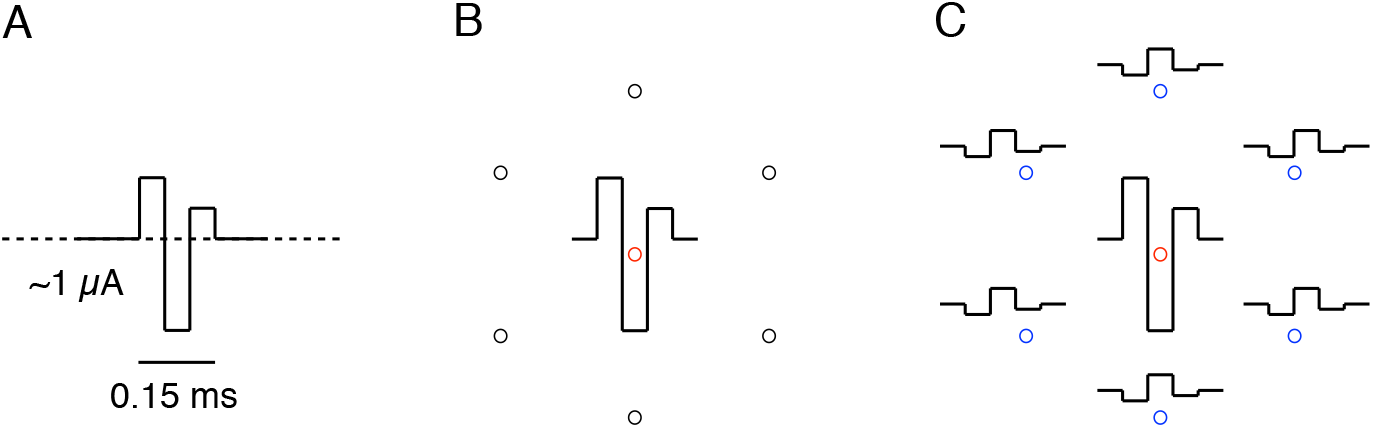
Schematic of distant return and local return stimulation. A: Anodic-first triphasic pulse passed through the central electrode. B: Center and neighbor electrodes (circles), and current waveform passed through the center electrode C: Electrodes and waveforms for local return stimulation. Current with ⅙ the amplitude of the central electrode current and opposite sign is passed through each of the 6 surrounding electrodes to create a local return. For B and C, the center stimulating electrode is outlined in red, active local return electrodes in blue, and inactive neighbor electrodes in black.

In both distant and local return stimulation, electrodes were stimulated one at a time, with ~8 ms between pulses. The order of stimuli was chosen pseudo-randomly, but was restricted so that each successive stimulating electrode was far from the previous and subsequent electrode. This was done to avoid stimulating the same cell(s) in rapid succession. 25 trials were completed for each pattern, at each of 39 current amplitudes (10% increments) between 0.1 and 3.7 μA.

### Visual stimulation and cell type classification

A dynamic white noise visual stimulus was used to characterize the light response properties of recorded RGCs. Spike sorting — the identification and segregation of spikes from different RGCs — in the presence of visual stimulation was accomplished using methods described previously (Litke et al., 2004; Field et al., 2007). RGC light responses were then summarized using the spike-triggered average (STA) stimulus obtained with white noise (Chichilnisky, 2001; Chichilnisky and Kalmar, 2002). STA properties and mosaic organization were used to uniquely identify the major RGC types (Field et al. 2007). The spatial extent of each STA (see Fig. 4) was summarized using an elliptical approximation of the receptive field spatial profile (Chichilnisky and Kalmar, 2002). In total 55 ON parasol, 26 OFF parasol, 3 ON midget, and 6 non-light responsive RGCs were analyzed. The non-light responsive cells were not definitively identified; however, their electrical images reveal that they are indeed RGCs (Greschner et al., 2014), and minority RGC types in the retina with large receptive fields may not respond as vigorously to the fine-grained white noise stimulation used as the major cell types.

### Electrical image and responses to electrical stimulation

The spikes recorded during electrical stimulation were analyzed with a custom semi-automated method (Jepson et al., 2013). Briefly, an automated algorithm separated spikes from the electrical artifact by grouping traces according to an artifact waveform estimate and the average recorded spike waveform from visual stimulation (see above). The electrical image, i.e. the average spatiotemporal pattern of voltage deflections produced on each electrode of the array during a spike (Litke et al., 2004), was calculated from visual stimulation data, and served as a template for the spike waveform of each cell to be detected during electrical stimulation. The electrically evoked RGC responses were visually inspected for sorting errors and manually corrected.

The positions of recorded cells were estimated using the electrical image (Litke et al., 2004)— specifically, the center of mass of the electrode with the largest amplitude recorded spike waveform and the two neighbors with largest amplitudes. RGCs were selected for analysis if they exhibited electrical images overlapping the stimulating electrode. These cells were then analyzed manually to confirm activation resulting from electrical stimulation. Finally, the activation threshold, defined as the current amplitude required to elicit spikes in 50% of the trials, was extracted. This process was repeated for all stimulating electrodes.

### Statistical analysis of threshold and selectivity changes

The relationship between local and distant returns was analyzed statistically using a resampling approach. To determine the estimated variation in the slopes of lines fitted to data in Fig. 2 and Fig. 3, points from the plots were resampled with replacement, producing simulated data sets with the same number of points as the real data, drawn from the distribution given by the measured data (Efron 1982). This resampling was performed repeatedly and, for each resampling, the slope of the least squares linear fit to the data was computed. Values in the text represent the slope obtained from the data, and 90% confidence intervals based on the distribution of slopes from the resampled data.

**Figure 2.**
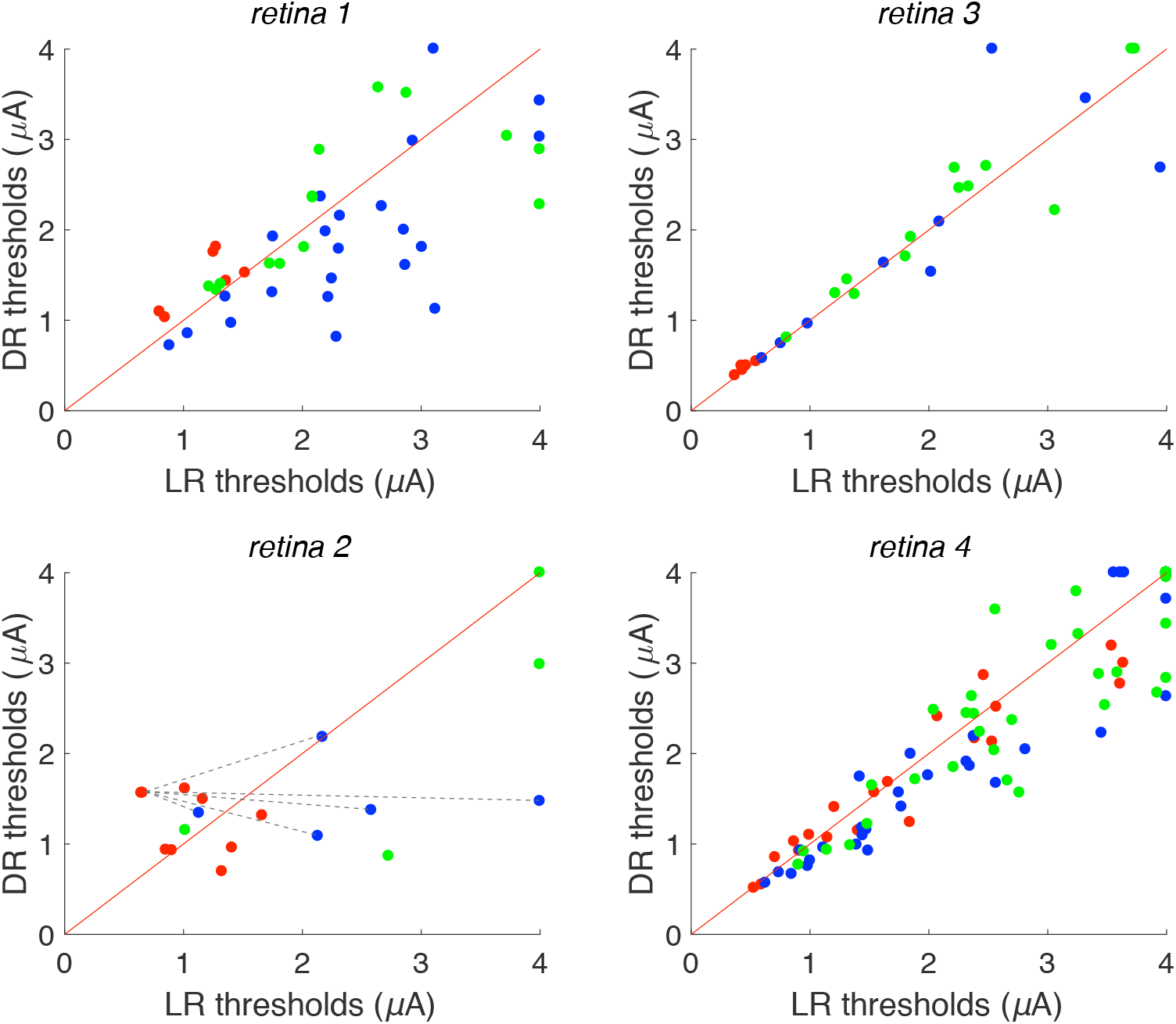
Local and distant return activation thresholds. Scatter plots for four different preparations show current threshold of analyzed cells to stimulation via distant return (DR) and local return (LR). Red diagonals denote the reference line y = x. Points along the right (top) edges of the plots indicate conditions in which no cellular activation was observed at the highest current level used with local return (distant return). Red points indicate near cells (<30 μm from stimulating electrode), blue points indicate intermediate cells (30-60 μm), green points indicate far cells (> 60 μm). An example of the method used to calculate predicted improvement in selectivity (see Fig. 3) is illustrated in retina 2: dashed lines show all possible pairings between a single red point and all blue points; similar pairings were made for all red points. Data are shown from 90 distinct cells, 199 distinct cell-electrode pairs.

**Figure 3.**
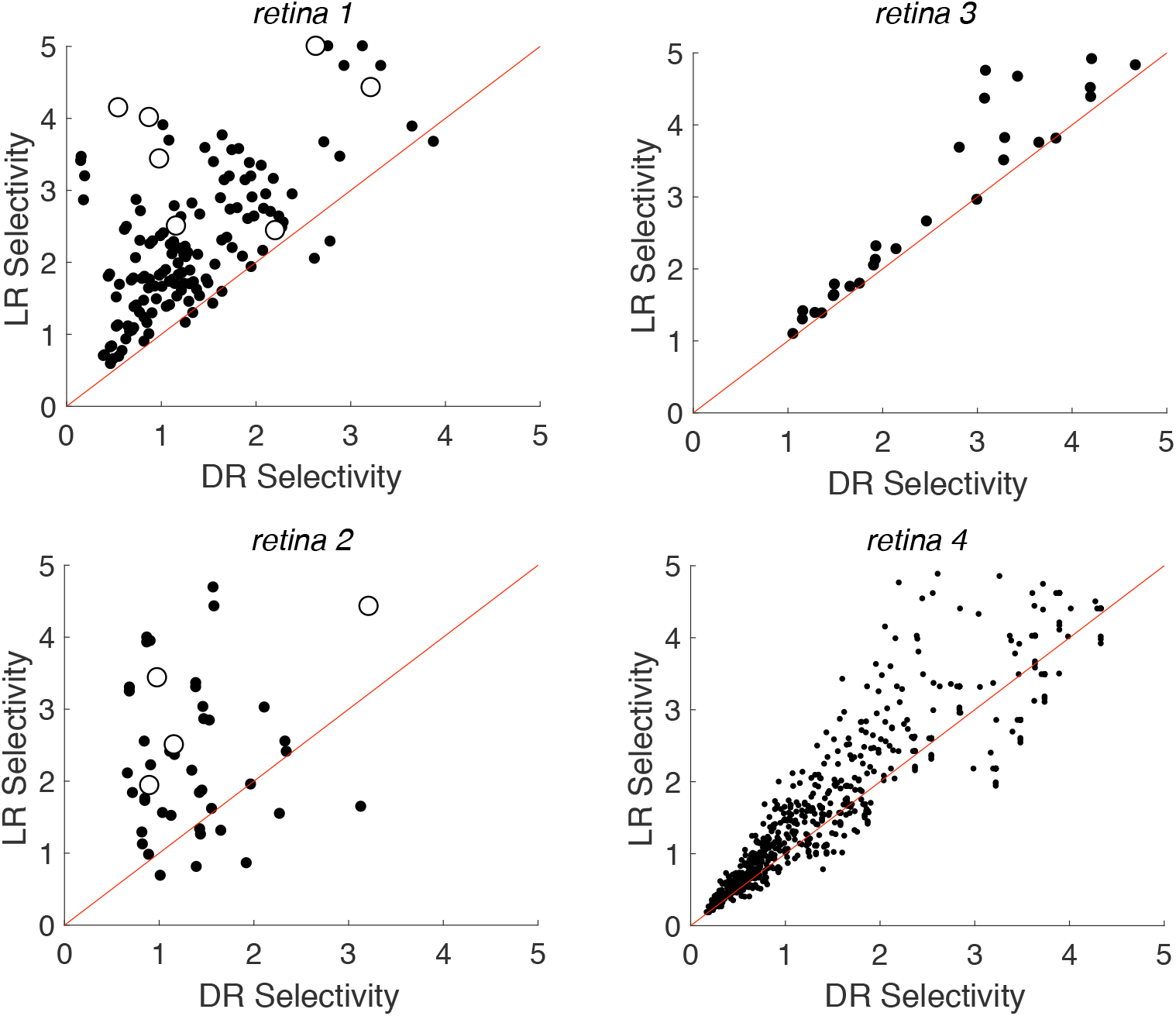
Impact of local returns on selectivity. Each point in the scatter plots indicate selectivity calculated for a near target cell (red points from Fig. 2) and an intermediate non-target cell (blue points from Fig. 2) with local return (LR) and distant return (DR) stimulation. Red diagonals denote the reference line y = x. Selectivity is defined as the ratio of non-target cell threshold to target cell threshold. Open symbols represent cases in which the target and non-target cells were actually recorded and stimulated simultaneously on the same set of electrodes

## Results

Electrical recording and stimulation were performed on isolated macaque monkey retina using large-scale high-density multi-electrode arrays (see Methods). Direct electrical activation of RGCs using weak current pulses (~1 μA; see Fig 1; Sekirnjak et al., 2006) passed through one or more electrodes was examined using different geometries of stimulation. To determine the effect of local return stimulation on the activation of individual RGCs, the results of stimulation through a central electrode with either a distant return (bath ground; ~1 cm) or local return (surrounding 6 electrodes; 60 μm) were compared (see Fig. 1). After performing repeated stimulation with brief current pulses (0.15 ms) at each of 39 current levels (0.1-3.7 μA), a current level that would produce RGC activation on 50% of trials was calculated (see Methods) and defined as the activation threshold. In all cases, stimulation was performed near the soma. This was ensured by using an electrode that recorded unambiguous somatic action potentials from the cell in response to light stimulation, and that produced a relationship between applied current and response probability less steep than is observed with axonal stimulation. To account for the geometry of cells with respect to the electrodes in the analysis that follows, analyzed cells were divided into three categories: *near* the stimulating electrode (<30 μm), *intermediate* distance (30-60 μm), and *far* from the stimulating electrode (>60 μm).

### Effect of local returns on RGC activation thresholds varies with distance and across recordings

Comparison of thresholds obtained with local and distant return stimulation yielded systematic trends within retinas, but somewhat variable trends across retinas (Fig. 2, Table 1). To summarize the relation between local and distant return stimulation for each retina and cell group, the slopes of the relation between local return and distant return thresholds (Fig. 2) was computed, along with 90% confidence intervals on the slope obtained by resampling (see Methods; Table 1). For cells near the stimulating electrode (red points in Fig. 2), local return stimulation produced significantly lower thresholds (slopes larger than 1) in two retinas (1,3), similar thresholds in one retina (2), and higher thresholds in one retina (4). For intermediate distance cells (blue points), local return produced systematically higher thresholds in three retinas (1,2,4) and similar thresholds in one retina (3). For far cells (green points), local return produced slightly higher thresholds in two retinas (2,4) and similar thresholds in two retinas (1,3). In summary, local return stimulation produced inconclusive results in near cells, but tended to produce higher thresholds in intermediate and perhaps far cells (Table 1).

**Table 1.** Change in thresholds with local vs. distant return. Cells are separated by cell group (near, intermediate, far). Each entry indicates the fitted slope of the data in Fig. 2, with 90% confidence intervals on the slope obtained by resampling from the data with replacement 1000 times (see Methods). The last row indicates the results of pooling the results from all cells across all four retinas.

| *LR vs. DR thresholds*<br>*mean (90% confidence interval)* |  |  |  |
| --- | --- | --- | --- |
| | near (<30 μm) | intermediate (30-60 μm) | far (>60 μm) |
| retina 1 | 1.22 (1.09, 1.37)<br>n = 6 | 0.698 (0.538, 0.840)<br>n = 23 | 0.926 (0.785, 1.09)<br>n = 14 |
| retina 2 | 1.04 (0.830, 1.32)<br>n = 9 | 0.572 (0.418, 0.860)<br>n = 5 | 0.757 (0.496, 1.00)<br>n = 4 |
| retina 3 | 1.05 (1.01, 1.10)<br>n = 5 | 0.916 (0.765, 1.13)<br>n = 10 | 1.03 (0.935, 1.09)<br>n = 13 |
| retina 4 | 0.916 (0.858, 0.981)<br>n = 20 | 0.929 (0.874, 0.974)<br>n = 42 | 0.955 (0.923, 0.985)<br>n = 48 |
| pooled | 0.953 (0.892, 1.02)<br>n = 40 | 0.845 (0.781, 0.907)<br>n = 80 | 0.971 (0.941, 0.998)<br>n = 79 |

This trend suggests that local returns could potentially be useful in focusing electrical activation near the stimulating electrode and avoiding the activation of cells at greater distances. However, the variability across cells and retinas, which could arise from several factors (see Discussion), was substantial. Given that such variability could occur in epiretinal prostheses as well, it is unclear from the above analysis alone how reliable and effective local returns may be in practice for stimulating individual cells while avoiding others.

### Local returns enhance selective activation of target cell over non-target cells

Therefore, to test the practical implications of these findings, a second analysis was performed, focused on whether local returns could be used to enhance selectivity of stimulation of a target cell over a nearby non-target cell. Specifically, a surrogate analysis was performed using the single-cell electrical stimulation data (Fig. 2), to test the situation in which one cell is targeted for stimulation near the stimulating electrode, while attempting to avoid activation of another cell at an intermediate distance. Note that, in reality, the pairs of cells analyzed were usually stimulated by different electrodes on the array (but see below), because the number of cells available for analysis was limited by the difficulty of reliably spike sorting data from many cells with each stimulating electrode. The surrogate pair analysis was therefore performed by taking all possible pairings between near cells (<30 μm from their stimulating electrode), considered to be the *target* cell, and intermediate cells (30-60 μm from their stimulating electrode), considered to be the *non-target* cell, within the same retina (see Fig. 2, retina 2 for illustration of pairing).

Selectivity, defined as the ratio of non-target cell threshold to target cell threshold, was computed for both local and distant return stimulation. High selectivity corresponds to more favorable circumstances for activation of the target cell without activation of the non-target cell. The selectivity for activating a near target cell over an non-target intermediate cell was systematically higher in the local return than the distant return configuration (Fig. 3). Slopes fitted to these data, along with 90% confidence intervals for the slopes obtained by resampling (Table 2), reveal a clear trend. Selectivity with local returns was substantially higher in two retinas (1,2), slightly higher in one retina (4), and similar in one retina (3). Selectivity was also higher in the data pooled across retinas. The results were similar when computing selectivity for a near cell relative to a non-target cell from the far category (Table 2).

**Table 2.** Change in selectivity with local vs. distant return. The first column indicates the fitted slope, and the confidence interval on the slope obtained by resampling, for local return vs. distant return selectivity, using cell pairs composed of one near cell and one intermediate cell (data from Fig. 3). The second column indicates the slopes for pairs composed of one near cell and one far cell. Details as in Table 1.

| *LR vs. DR selectivity measured by thresholds*<br>*mean (90% confidence interval)* |  |  |
| --- | --- | --- |
|  | near vs. intermediate | near vs. far |
| retina 1 | 1.47 (1.39, 1.56) | 1.37 (1.31, 1.43) |
| retina 2 | 1.57 (1.28, 1.91) | 1.32 (1.12, 1.55) |
| retina 3 | 1.09 (0.980, 1.20) | 1.04 (0.997, 1.10) |
| retina 4 | 1.03 (1.01, 1.05) | 1.02 (1.01, 1.03) |
| pooled | 1.29 (1.01, 1.73) | 1.10 (0.996, 1.34) |

In addition, although the above results were mostly obtained with surrogate cell pairs pooled across stimulating electrodes, in a small number of cases, the target and non-target cells were actually stimulated by the same electrode (Fig. 3, open symbols) – in other words, the comparative activation of the two cells was real rather than simulated, providing a more direct empirical test of the effectiveness of local returns. In all 11 cell pairs of this kind (retinas 1 and 2) local return stimulation produced substantially higher selectivity than distant return (p<0.001), confirming the results of surrogate analysis.

Selectivity was also evaluated using an alternative measure: the largest difference in activation probabilities, across the range of tested currents, between the target cell and the non-target cell. This measure provides an indication of the practical impact in terms of how frequently the target cell can be activated without the non-target cell. By this measure, selectivity was also enhanced in local return relative to distant return stimulation. Specifically, in the four retinas tested, mean selectivity across cells was substantially higher for local returns in two retinas (1,2), slightly higher in one retina (4), and similar in one retina (3) (Table 3), the same trend that was observed with the selectivity measure based on stimulation threshold. The increases in the activation probability of the target cell were in some cases substantial (e.g. ~50% in retina 1). The effect was slightly weaker when computing selectivity for a target near cell relative to a non-target far cell (Table 3) rather than a non-target intermediate cell. Also, the median (rather than mean) effects were weaker using this measure of selectivity, suggesting that a minority of cells with large increases dominated the results obtained with the mean (not shown). Finally, another measure, the largest ratio of activation probabilities across the range of stimulation currents, yielded results similar to those obtained with the original threshold measurement (not shown).

**Table 3.**
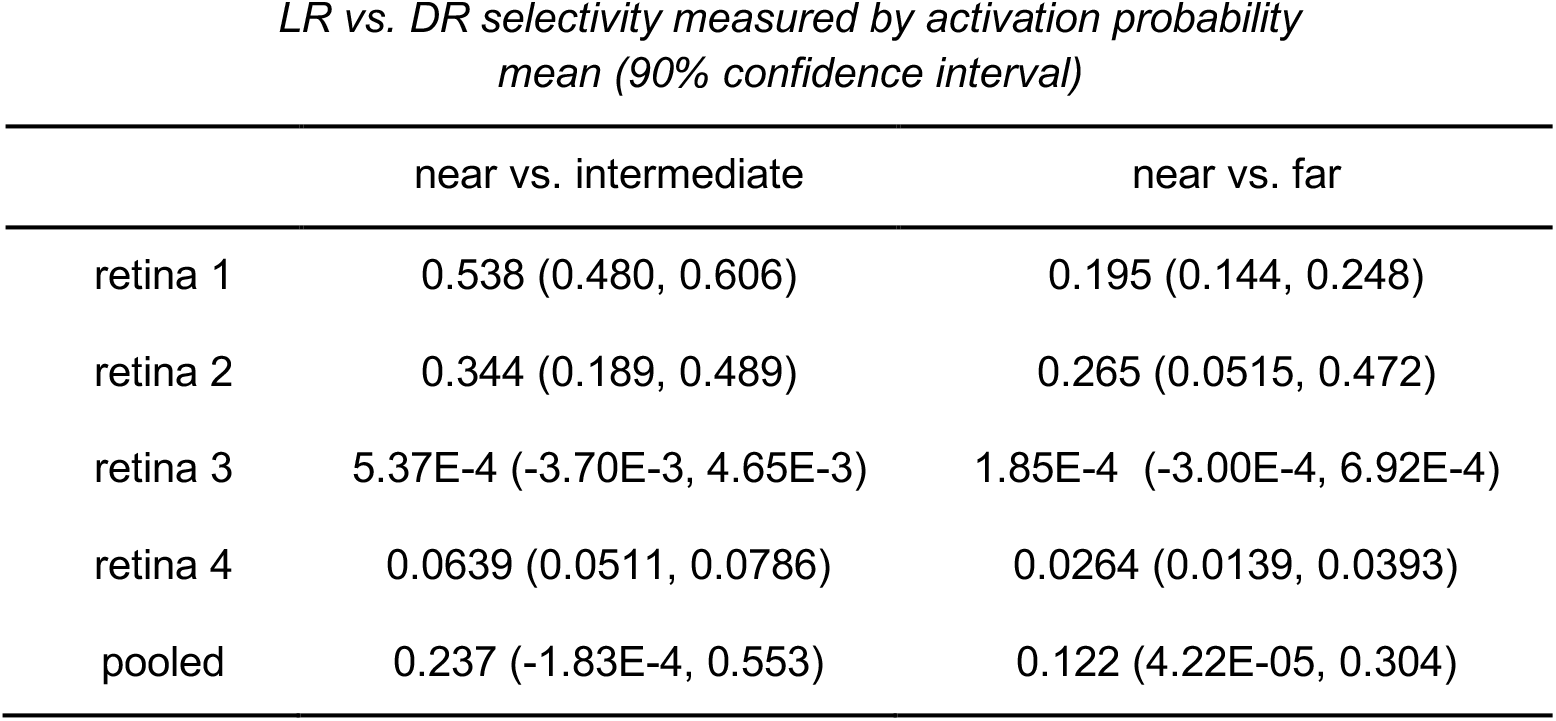
Change in selectivity with local vs. distant return, using alternate selectivity measure. Selectivity was defined as the largest difference, across current levels, in the activation probabilities of target and non-target cells. Selectivity was computed for each cell pair in the local return condition, and separately in the distant return condition. The table shows the difference in these two selectivity values. Each entry shows the mean and 90% confidence interval (obtained by resampling), across target/non-target cell pairs. Positive values indicate increases in selectivity with local return stimulation, negative values indicate decreases. The cell pairs are segregated into near vs. intermediate and near vs. far, based on distance from the stimulating electrode (see Results).

### Selective activation of different RGC types encoding adjacent/overlapping spatial locations

The cell pairs recorded on the same electrodes in the above analysis also revealed the cellular spatial resolution of increases in selectivity. Specifically, in five cases, the target and non-target cells were of opposite sign (ON and OFF parasol) and had immediately adjacent or overlapping receptive fields within the mosaics of these two cell types (Fig. 4, left). Thus, local return stimulation made it possible to more selectively stimulate two cells in very close proximity in terms of the portion of the visual field they encoded. Furthermore, it enhanced the ability to avoid indiscriminate activation of cells with opposite light response polarities, which could be important for artificial vision (see Discussion). Finally, the threshold and activation probability changes caused by local return stimulation were substantial and directly visible in the activation curves (Fig. 4, right).

**Figure 4.**
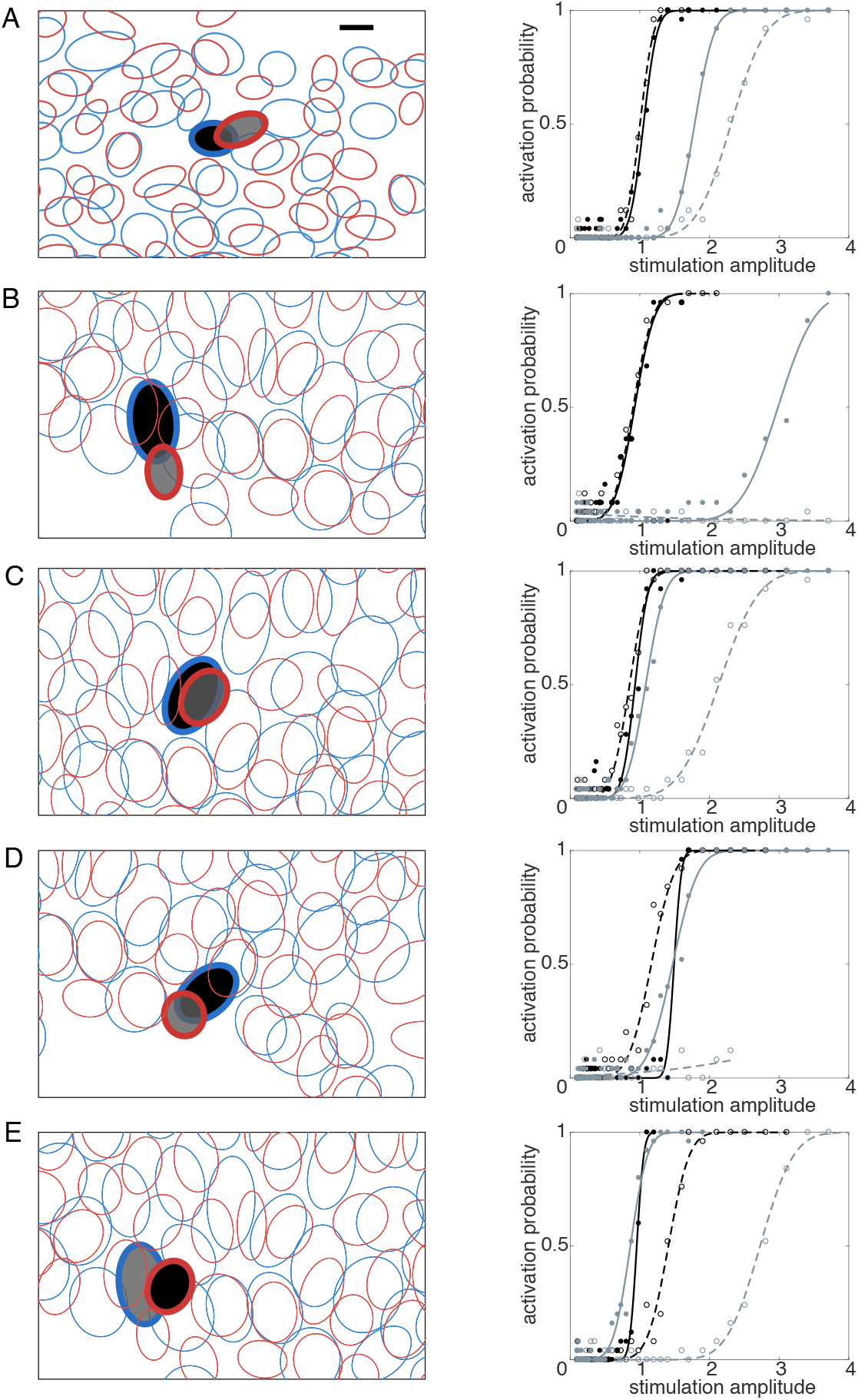
Relative locations and activation curves for target and non-target cells exhibiting selectivity enhancement. Each panel in the left column shows the mosaic of receptive fields (ellipses) of recorded ON (blue) and OFF (red) parasol cells over a region of a particular recording, measured using reverse correlation with white noise and fitted with elliptical Gaussians (see Methods). Cell pair A was from retina 1, cell pairs B-E from retina 2. Filled ellipses with bold outlines indicate target (black) and non-target (gray) cell pairs, stimulated and recorded with the same electrodes. The stimulating electrode used was different for each cell pair. Note that the target cell in B was also the non-target cell in E. The response curves for each pair are shown in the right column. Solid lines indicate response to distant return stimulation; dashed lines indicate the response to local return stimulation. In each case, local return stimulation shifted the non-target (gray) activation probability curve to the right, resulting in a higher activation threshold, while affecting the target cell (black) activation curve less or not at all. In B and D, with local return, no activation of the non-target cell was observed over the entire stimulation range. Scale bar: 100 μm.

## Discussion

The results indicate that local returns can enhance the selectivity of epiretinal stimulation of RGCs, at cellular resolution, in the primate retina. This finding is broadly consistent with the well-established fact that local return stimulation can be used to restrict the electrical activation of neurons by focusing the electric field (Abramian et al., 2011; Habib et al., 2013). The primary novel aspect of the present findings is that increases in resolution occur over very small spatial scales that enhance the probability of single-cell activation. Also, the results show that the selectivity enhancement with local returns is systematic, in spite of variable effects on individual cell thresholds. Finally, the results show that cellular resolution activation can be used to differentially target ON and OFF cells encoding immediately adjacent or overlapping areas of the visual field.

On a practical level, these findings suggest that a retinal prosthesis designed to operate at single-cell resolution could be improved by the use of local returns, to selectively target cells near each stimulating electrode while avoiding the activation of neighboring cells. To exploit this higher selectivity, an ideal device would record electrically evoked activity to determine the optimal current amplitude for activating the target cell above threshold, while remaining below threshold for the non-target cell(s). Also, it would be important for such a device to identify the degree to which passing axons are activated, a major issue for existing epiretinal devices (Nanduri, 2011; Weitz et al., 2015), but one that is likely to be manageable in a subset of cells if the device is able to record and calibrate stimulation current levels (Grosberg et al., 2017). In the present study, only somatic activation was evaluated. This was accomplished by using electrodes that recorded unambiguous somatic spikes, and that also produced activation curves less steep than those normally observed with axonal activation. A thorough examination of axon activation would require further experiments.

The cellular resolution of the increases in selectivity with local returns was revealed by the fact that a target cell could be more effectively activated relative to a non-target cell encoding an immediately neighboring or overlapping location in the visual field (Fig. 4). Thus, local returns can effectively enhance the highest possible visual resolution. Furthermore, local returns permitted activation of an ON cell more effectively than an immediately adjacent or overlapping OFF cell, or vice-versa. This is significant because the indiscriminate activation of ON and OFF cells encoding the same location in the visual field is a salient example of how poor selectivity can cause conflicting visual information to be transmitted to the brain (see Goetz and Palanker, 2016).

The substantial variability of the effects of local returns across cell pairs and retinas could have several origins. One factor is the more restricted electric field in the depth dimension of the retina with local returns (Flores et al., 2016), the effects of which may depend on features that are difficult to measure and different in different retinas, such as the thickness of the inner limiting membrane and axon fiber layer, and the degree to which the retina is pressed against the electrode array. Another factor is the non-radially-symmetric electric field produced by the six surrounding electrodes forming the local return, both within the ring of six electrodes, and outside it, which would vary with the locations of the target and non-target cells relative to the electrodes. A third contributing factor is errors in estimating the position of RGCs relative to the electrodes. Given that these factors are difficult to estimate, and were not estimated in the current work, their contributions to variability across cell pairs and retinas are unknown. Furthermore, it remains unclear whether these factors could produce the weaker improvements in selectivity for far non-target cells, compared to intermediate non-target cells. Although all of these factors deserve further examination, the overall trend across the experimental variables examined was that local returns enhanced selectivity significantly.

The majority of cells examined in this study were ON and OFF parasol cells, two of the major RGC types in the primate retina, comprising ~16% of the RGC population (see Dacey, 2004for review). A few ON midget cells were also examined; these cells comprise ~25% of all RGCs. Analysis focused primarily on parasol cells because their large spikes were easier to sort reliably in the presence of electrical artifacts. For a more comprehensive view of how local returns can impact selectivity of RGC stimulation, it may be necessary to more thoroughly explore the properties of cell types other than parasol cells, which would require further experimentation. An additional challenge is that manually spike sorting voltage traces recorded after electrical stimulation is arduous and time-consuming (see Jepson et al., 2012; Grosberg et al., 2017). One potential solution would be the development of algorithms to automatically sort spikes in these data, an effort that is underway (see Mena et al., 2017).

The electric field in the region near the stimulating electrode is predicted to be inhomogeneous in the present experiments, because the local return consisted of six individual electrodes surrounding the stimulating electrode, rather than a uniform ring. This inhomogeneity could make it more difficult to activate cells in certain regions near the stimulating electrode. The results obtained from the four retinal preparations generally support this possibility: although intermediate distance cells usually had higher local return thresholds, there were exceptions. It is possible that a surrounding return ring electrode would be more effective. However, the simple triangular lattice electrode arrangement has the advantage of flexibility: in a clinical device, such an arrangement could be used to provide a higher density of possible stimulation locations, while at the same time allowing some of the advantages of local return stimulation.

## Acknowledgements

This work was supported by National Eye Institute grants EY021271 (EJC) and P30 EY019005 (EJC) and F32EY025120 (LG), Stanford Neurosciences Institute (EJC), Stanford UAR Major Grant (VF), Stanford Bio-X USRP Fellowship (VF), Polish National Science Centre Grant DEC-2013/10/M/NZ4/00268 (PH). AGH UST, task No. 11.11.220.01/4 within subsidy of the Ministry of Science and Higher Education (WD), Pew Charitable Trusts Scholarship in the Biomedical Sciences (AS). We thank Bill Newsome, Tirin Moore, Adrienne Mueller, Jose Carmena, and Christie Ferrecchia for access to primate retinas, and Daniel Palanker for useful discussions.

